## Supplementary figures and images for "Diverse prey capture strategies in teleost larvae"

### Suppl. Fig. 1

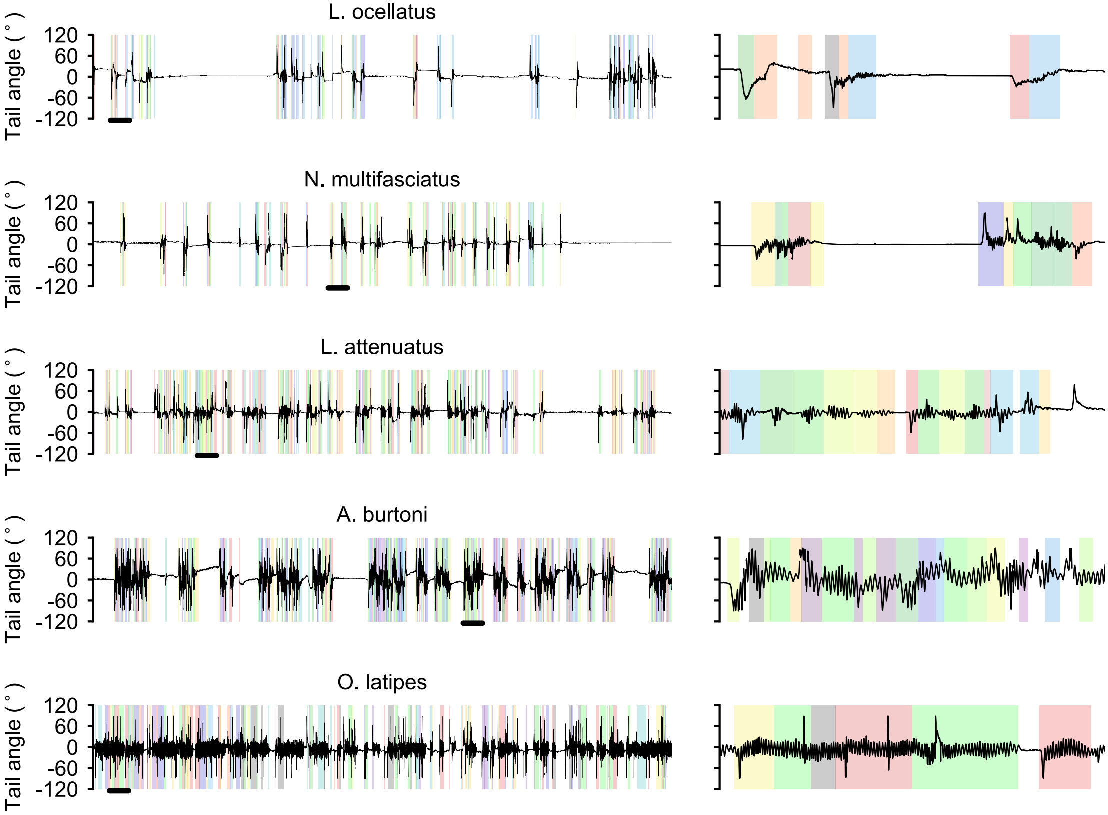

### Suppl. Fig. 2

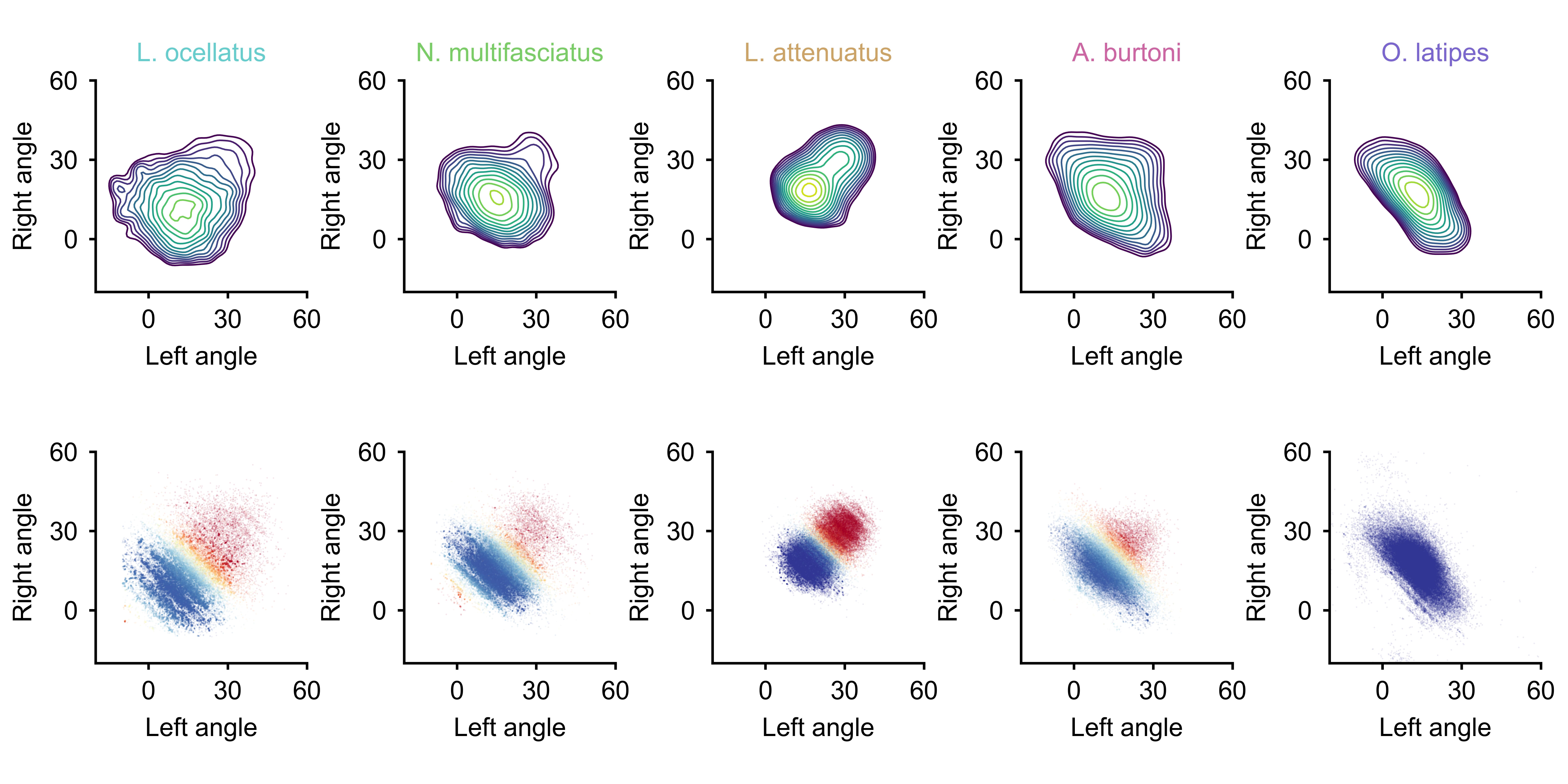

### Suppl. Fig. 3

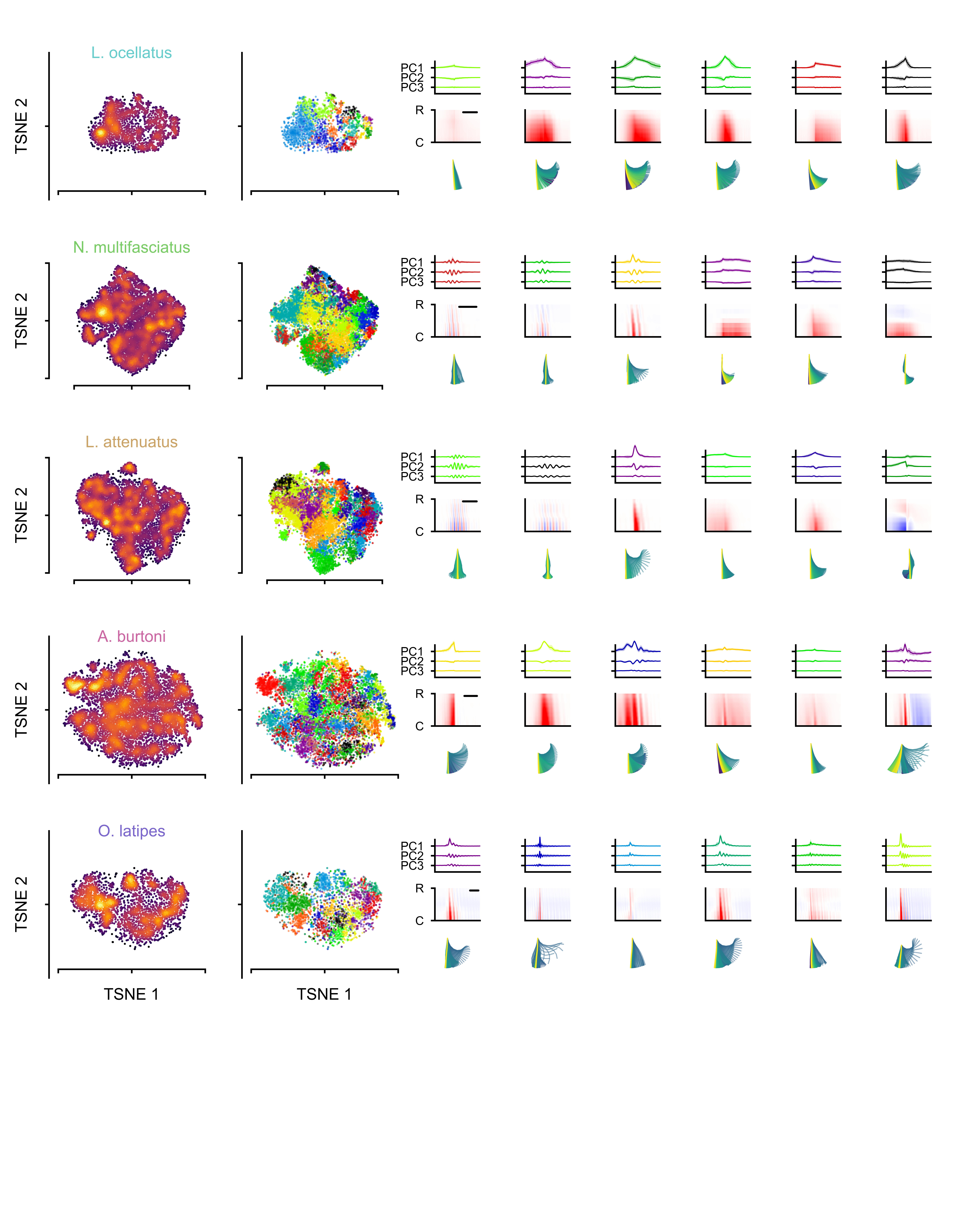
